# Poly(A) Binding Protein Nuclear 1 regulates the polyadenylation of key synaptic plasticity genes and plays a role in homeostatic plasticity

**DOI:** 10.1101/121194

**Authors:** Irena Vlatkovic, Sivakumar Sambandan, Georgi Tushev, Mantian Wang, Irina Epstein, Caspar Glock, Nicole Fuerst, Iván Cajigas, Erin Schuman

**Affiliations:** Max Planck Institute for Brain Research, Max-von-Laue Str. 4, 60438 Frankfurt.

**Keywords:** polyadenylation, PABPN1, hippocampal neurons, homeostatic plasticity, Camk2a, Gria2, 3’ untranslated regions

## Abstract

Polyadenylation is a nuclear process that involves the endonucleolytic cleavage of RNA transcripts and the addition of poly(A) tails. The cleavage often takes place at different positions within the same RNA transcript, generating alternative 3’ends. Polyadenylation regulates mRNA localization, stability and translation and is likely to regulate complex processes such as synapse formation, synaptic plasticity and memory. Here we examined whether PolyA Binding Protein Nuclear 1 (Pabpn1), an RNA binding protein known to regulate alternative polyadenylation and polyA tail length in other systems, regulates neuronal mRNA function. Using immunocytochemistry we determined that Pabpn1 is present in both hippocampal slices and cultured hippocampal neurons. Applying shRNAs to knock-down Pabpn1 we discovered that Pabpn1 regulates the mRNA abundance and localization of key synaptic plasticity genes including Calcium/Calmodulin Dependent Protein Kinase II Alpha (Camk2a) and Glutamate Ionotropic Receptor AMPA Type Subunit 2 (Gria2). Furthermore, Pabpn1 knock-down prevented the homeostatic scaling of synaptic transmission elicited by bicuculline. These data demonstrate a link between Pabpn1, polyadenylation and neuronal plasticity.

## Introduction

Neurons are distinct with their complex morphology and structural and functional compartmentalization. Recent studies have highlighted the importance of the decentralized translational machinery and how it underlies important cellular processes such as synapse formation, plasticity and memory (Holt and Schuman, 2013). The localization of mRNA and its regulated translation are integral components of local protein synthesis in neurons. Like in non-neuronal systems, the 3’ untranslated region (3’UTR) as well as the polyA tail length of an mRNA molecule harbor information and regulatory nodes for compartment targeting, translational efficiency and stability (Martin and Ephrussi, 2009; Weill et al., 2012). Recently, unusually long 3’UTRs were observed in mouse and human brain samples and many of these 3’UTRs belong to mRNAs that are alternatively polyadenylated (Miura et al., 2013; Taliaferro et al., 2016).

Polyadenylation takes place in the nucleus and involves the recognition of pre-mRNA sequences by a specific set of molecules (see (Elkon et al., 2013)). Besides the core polyadenylation proteins (e.g. Cleavage and Polyadenylation Specificity Factor, CPSF, and Cleavage Stimulation Factor, CSTF), proteins that can affect the choice of the mature 3’UTR isoform have been described. While Pabpn1 (Poly(A) Binding Nuclear Protein 1) was initially described as a protein that stimulates Poly(A) polymerase (PAP) during pre-mRNA polyadenylation and controls the polyA tail length, it has recently been associated with long non-coding RNA turnover, the hyperadenylation and decay of specific nuclear RNAs and the regulation of 3’UTR isoform choice (Beaulieu et al., 2012; Bresson and Conrad, 2013; de Klerk et al., 2012; Jenal et al., 2012; Kerwitz et al., 2003; Kuhn et al., 2009).

A specific mutation in Pabpn1 gives rise to oculopharyngeal muscular dystrophy (OPMD) (Brais et al., 1998; Chartier et al., 2015). In muscle cells, Pabpn1 promotes the preferential maturation of long 3’UTR isoforms (de Klerk et al., 2012; Jenal et al., 2012). To test for a potential role of Pabpn1 in 3’UTR lengthening in neurons, we first examined the abundance of Pabpn1 in the cortex, cerebellum, and hippocampus. Using an shRNA strategy, we discovered that knock down of Pabpn1 (Pabpn1-KD) leads to a decrease in the long 3’UTRs of several key synaptic plasticity genes. Unexpectedly, we found that the coding sequences of these genes are also decreased, suggesting that Pabpn1 influences the PAP efficiency, polyA tail length and likely the lifetime of the examined plasticity genes. Finally, we observed that Pabpn1-KD also blocks homeostatic scaling, a form of synaptic plasticity that requires mRNA translation (Schanzenbacher et al., 2016).

## Results

### Pabpn1 regulates polyadenylation of key synaptic plasticity genes

Pabpn1 is a ubiquitously expressed protein. To examine its expression pattern in different brain areas, we conducted immunostaining using an anti-Pabpn1 antibody. In the rat cortex, cerebellum and hippocampus Pabpn1 is prominently expressed in neuronal nuclei (Figure 1A). We additionally examined the localization of both the Pabpn1 mRNA and protein in cultured rat hippocampal neurons. We observed abundant Pabpn1 mRNA particles in the nuclei and cytoplasm (Figure 1B). The nuclear Pabpn1 protein staining appeared as granular structures, typically covering about 50% of the nuclear volume (Figure 1B).

**Figure 1.**
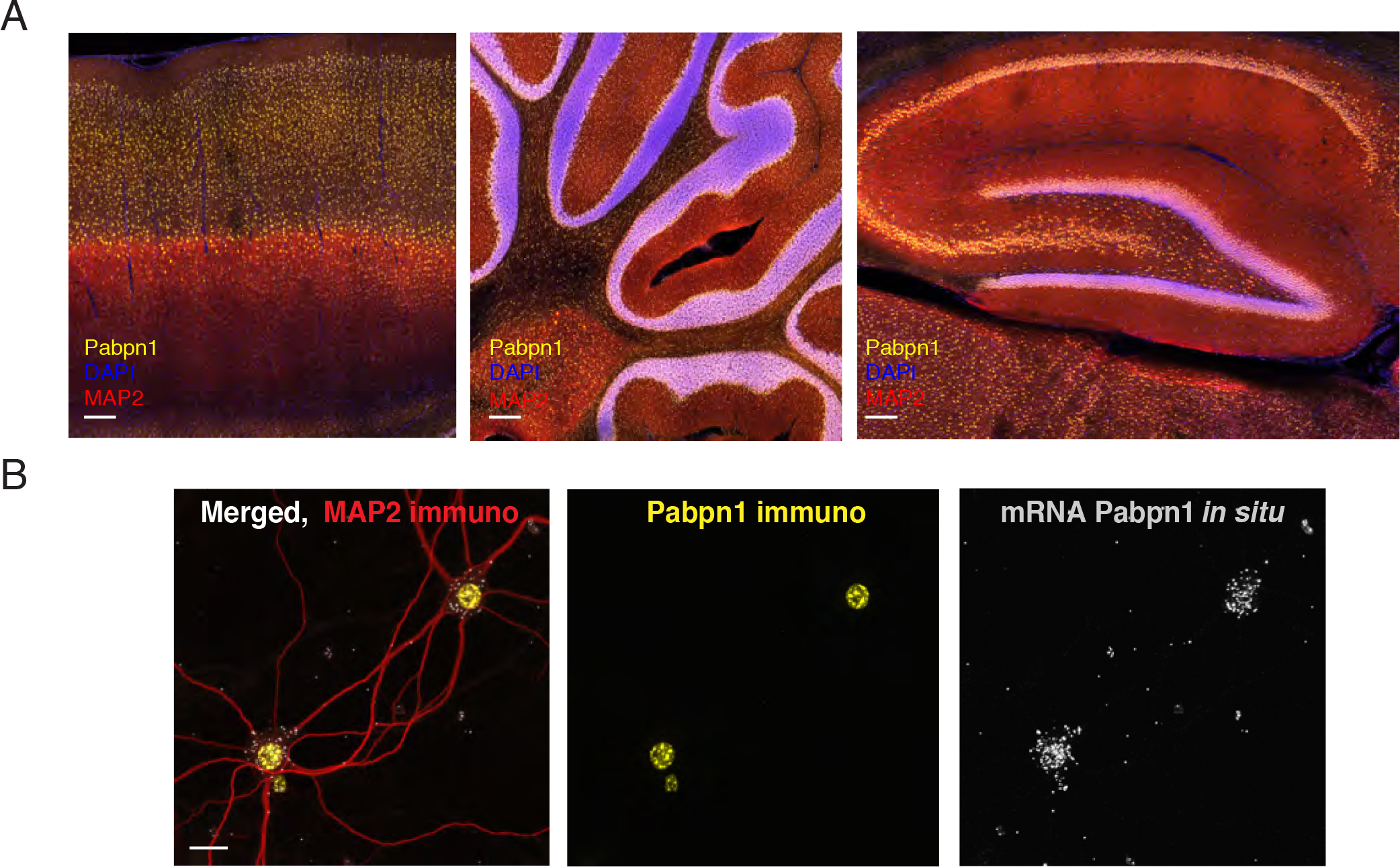
Pabpn1 is abundant in hippocampal neurons. A. Immunostaining of Pabpn1 (yellow) in rat cortex, cerebellum and hippocampus. Scale= 200 *µ*m. MAP2 (red) stains neurons and DAPI (blue) nuclei. B. Representative images of merge, immunostaining of Pabpn1 (yellow) and *in situ* hybridization of Pabpn1 (grey particles) in rat hippocampal neurons. MAP2 (red) stains neurons. Scale = 10*µ*m.

In order to identify potential Pabpn1 mRNA targets, we bioinformatically analyzed a rat primary hippocampal neuron 3’end deep sequencing dataset (Tushev et al, unpublished). We searched for all mRNAs that are predicted targets of Pabpn1 using the following criterium: mRNAs which possess at least two 3’UTR isoforms of which the shorter isoform ends with a less common poly(A) signal (non- canonical) while the extended 3’UTR isoform ends with the most common polyA site (canonical, AAUAAA; PAS) (Jenal et al., 2012). Using these criterium, we detected >1200 candidate Pabpn1 target RNAs, (data not shown). Notably, among the predicted targets we found a set of genes functionally important for synaptic plasticity including Calcium/Calmodulin Dependent Protein Kinase II Alpha (Camk2a), the Glutamate Ionotropic Receptor AMPA Type Subunit 2 (Gria2) and the Glutamate Ionotropic Receptor NMDA Type Subunit 2A (Grin2a) (Figure 2A).

**Figure 2.**
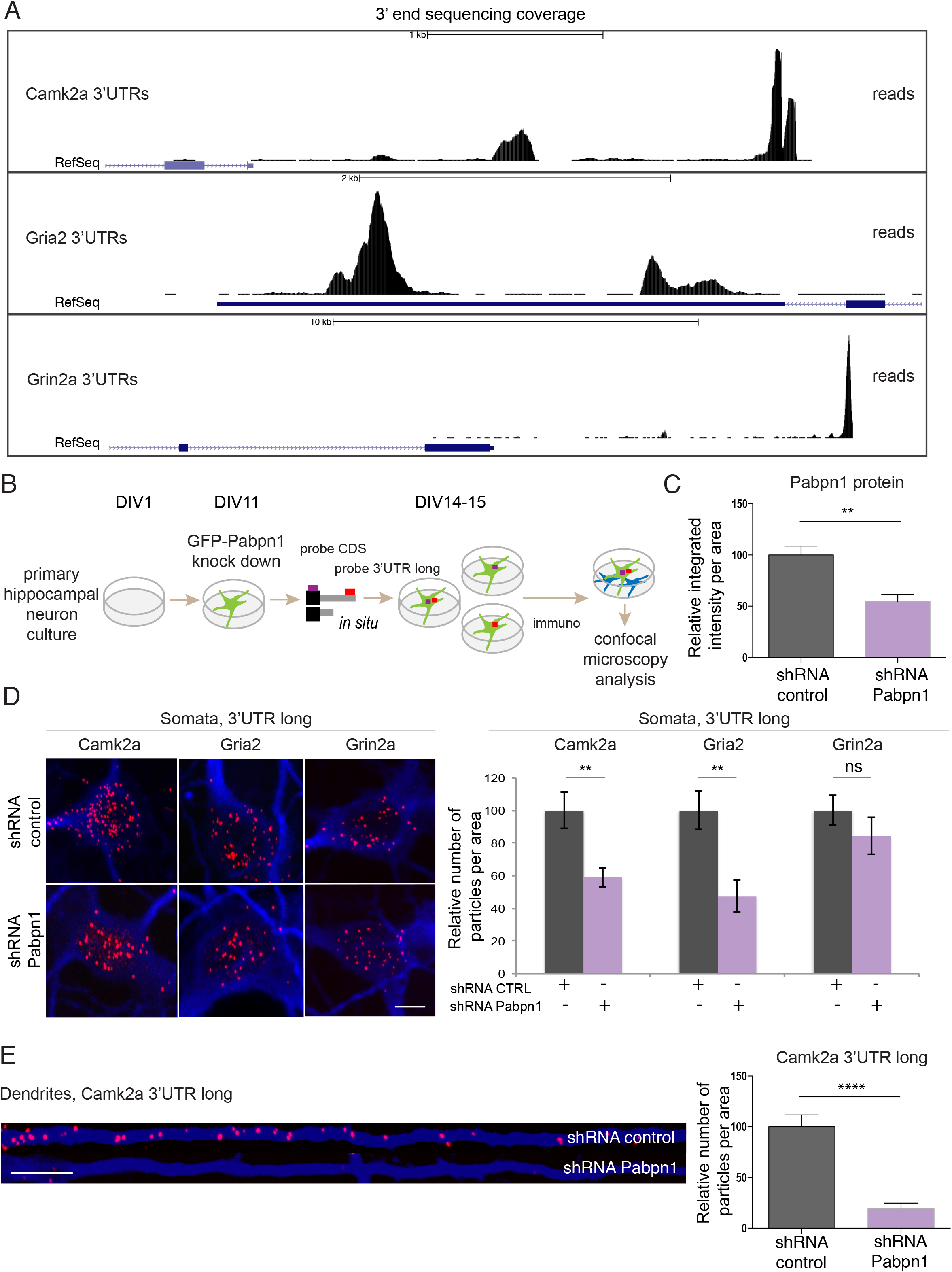
Pabpn1 knock down downregulates long 3’UTR isoforms of Camk2a and Gria2. A. Visualization of 3’end sequencing coverage in primary hippocampal neuron culture for Camk2a 3’UTRs, Gria2 3’UTRs and Grin2a 3’UTRs. UCSC web browser screenshots of rn5 display from the top to the bottom: scale, 3’end sequencing data in black and RefSeq annotation in dark blue. Black peaks of 3’end data represent distinct 3’UTR isoform ends. B. Illustration of the method including Pabpn1-KD, *in situ* hybridization and immunostaining of primary hippocampal neuron culture. C. Quantification of Pabpn1 immunostaining shows a decrease upon treatment with an shRNA against Pabpn1 (n= 9) compared to shRNA control neurons (n= 10), **p= 0.0015 D. Left panel: representative images of neurons treated with shRNA control and shRNA against Pabpn1 somata for Camk2a, Gria2 and Grin2a 3’UTR long (red particles), MAP2 (blue). Scale= 10 *µ*m. Right panel, from left to right: quantification of *in situ* hybridization data for long 3’UTRs of Camk2a, Gria2 and Grin2a shows a significant decrease in the relative number of particles per area in neurons treated with the shRNA against Pabpn1 for Camk2a and Gria2 3’UTRs (**p= 0.0011 and **p= 0.0026), while for Grin2a 3’UTR the change is not significant (n.s.), when compared to shRNA control samples normalized to 100. Numbers of analyzed neurons, from left to right: n= 21, 20, 14, 12, 32, 25. Whiskers represent Standard error of the mean (S.E.M). E. Left panel: Representative image of straightened dendrites. MAP2 stained dendrites are shown in blue, *in situ* hybridization of Camk2a 3’UTR long is shown in red. Right panel: quantification of Camk2a 3’UTR long particles per dendritic area shows a decrease upon Pabpn1-KD (****, p<0.0001, n=19, 19). Whiskers represent S.E.M.

To investigate a potential role of Pabpn1 in 3’UTR lengthening in hippocampal neurons, we used RNA interference, treating dissociated hippocampal cultured neurons with either a non-targeting shRNA control or an shRNA against Pabpn1 (Figure 2B). Neurons treated with the Pabpn1 shRNA exhibited a ∼ 50% knock down of the Pabpn1 protein (Figure 2C). To test if Pabpn1 plays a role in 3’UTR choice of the above predicted targets, we used an *in situ* hybridization approach with probes recognizing unique sequences of the long 3’UTR for the Camk2a, Gria2 or Grin2a mRNA. We analyzed the signal obtained from confocal imaging of *in situ* experiments and normalized the signal to the somatic or dendritic compartment area obtained by immunostaining (using an antibody against Map2). Pabpn1-KD led to a significant decrease in the long 3’UTR isoforms of Camk2a and Gria2 mRNA, while the long 3’UTR of the Grin2a mRNA was not affected (Figure 2D). Interestingly, the observed decrease of the long Camk2a 3’UTR isoform was more pronounced when we analyzed the expression in dendrites which exhibited an ∼80% decrease in signal, whereas the soma exhibited a ∼40% decrease (Figure 2E).

To test if the reduction in long 3’UTR mRNA is caused by an effect of Pabpn1 on alternative polyadenylation or polyA tail lengthening, we performed *in situ* hybridization using probes recognizing the coding sequences of Camk2a, Gria2 or Grin2a. If alternative polyadenylation is the reason for the observed decrease in Camk2 and Gria2 long 3’UTR isoforms, no change in the coding sequence (CDS) is expected, since in the simplest scenario, a decrease in the levels of long 3’UTR isoforms should result in an increase in the levels of the short 3’UTR isoforms and hence no net effect on the total transcript level (Jenal et al., 2012). When *in situ* probes directed against the coding sequence were used, both the Gria2 and the Camk2a transcript population showed decreases in Pabpn1 KD neurons. In the case of Gria2, the CDS probes showed a similar magnitude decrease as observed for the 3’UTR probe. The Camk2a CDS showed an even more pronounced decrease when compared to the 3’UTR probe (∼80% decrease in the soma and 80% - 90% decrease in dendrites) (Figure 3A and B). These data suggest that the effects of Pabpn1 knock-down cannot be attributed to a simple change in the choice of polyadenylation site, but rather may affect polyadenylation in general and possibly the lifetime of the mRNA.

**Figure 3.**
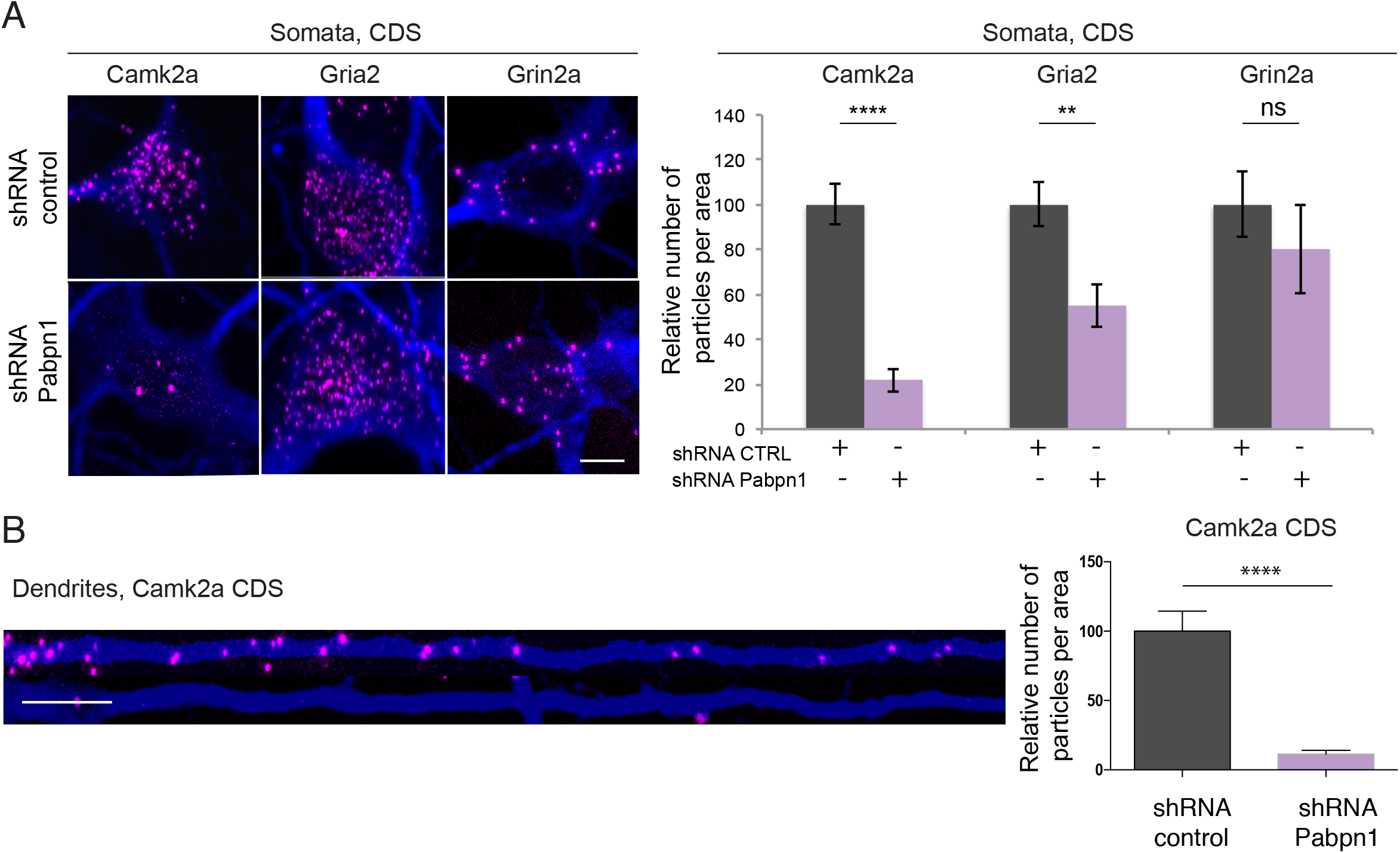
Pabpn1 knock down decreases the levels of Camk2a and Gria2 coding sequences (CDSs). A. Left panel: representative images of shRNA control and shRNA against Pabpn1 somata for Camk2a, Gria2 and Grin2a coding sequence (purple particles), MAP2 (blue). Scale= 10 *µ*m. Right panel, from left to right: quantification of *in situ* hybridization data for Camk2a, Gria2 and Grin2a shows a decrease in shRNA against Pabpn1 for Camk2a and Gria2 (****p<0.0001 and **p=0.0028). Grin2a is not changed significantly (n.s.). Numbers of analyzed neurons, from left to right: n= 21, 19, 14, 14, 21, 15. Whiskers represent S.E.M. B. Left panel: representative image of straightened dendrites. MAP2 stained dendrites are shown in blue, *in situ* hybridization of Camk2a coding sequence is shown in purple. Right panel: quantification of Camk2a particles per dendritic area shows a decrease upon Pabpn1-KD (****, p<0.0001, n=21, 19). Whiskers represent S.E.M.

### Pabpn1 knock down downregulates newly synthesized Camk2a protein

To examine whether the Pabpn1-KD elicited decrease in Camk2a mRNA affects Camk2a protein levels, we used the Puromycin-Proximity Ligation Assay (Puro-PLA), (tom Dieck et al., 2015). Following 3 days of Pabpn1-KD, we incorporated puromycin (4 min), a tRNA analog that leads to translation termination, into newly synthesized Camk2a. We then used puromycin- and Camk2a- specific antibodies and secondary antibodies coupled to oligonucleotides that allow rolling-circle amplification, to coincidently detect puromycin and Camk2a in the same molecule (tom Dieck et al., 2015) (Figure 4A). This allowed for an *in situ* visualization of the quantity of newly synthesized Camk2a protein upon Pabpn1-KD. In control experiments, the protein synthesis inhibitor anisomycin (40 *µ*M) was added 30 min prior to shRNA transfection. Using this strategy, we observed a significant decrease in newly synthesized Camk2a in both somata and dendrites following Pabpn1-KD (Figure 4B and C).

**Figure 4.**
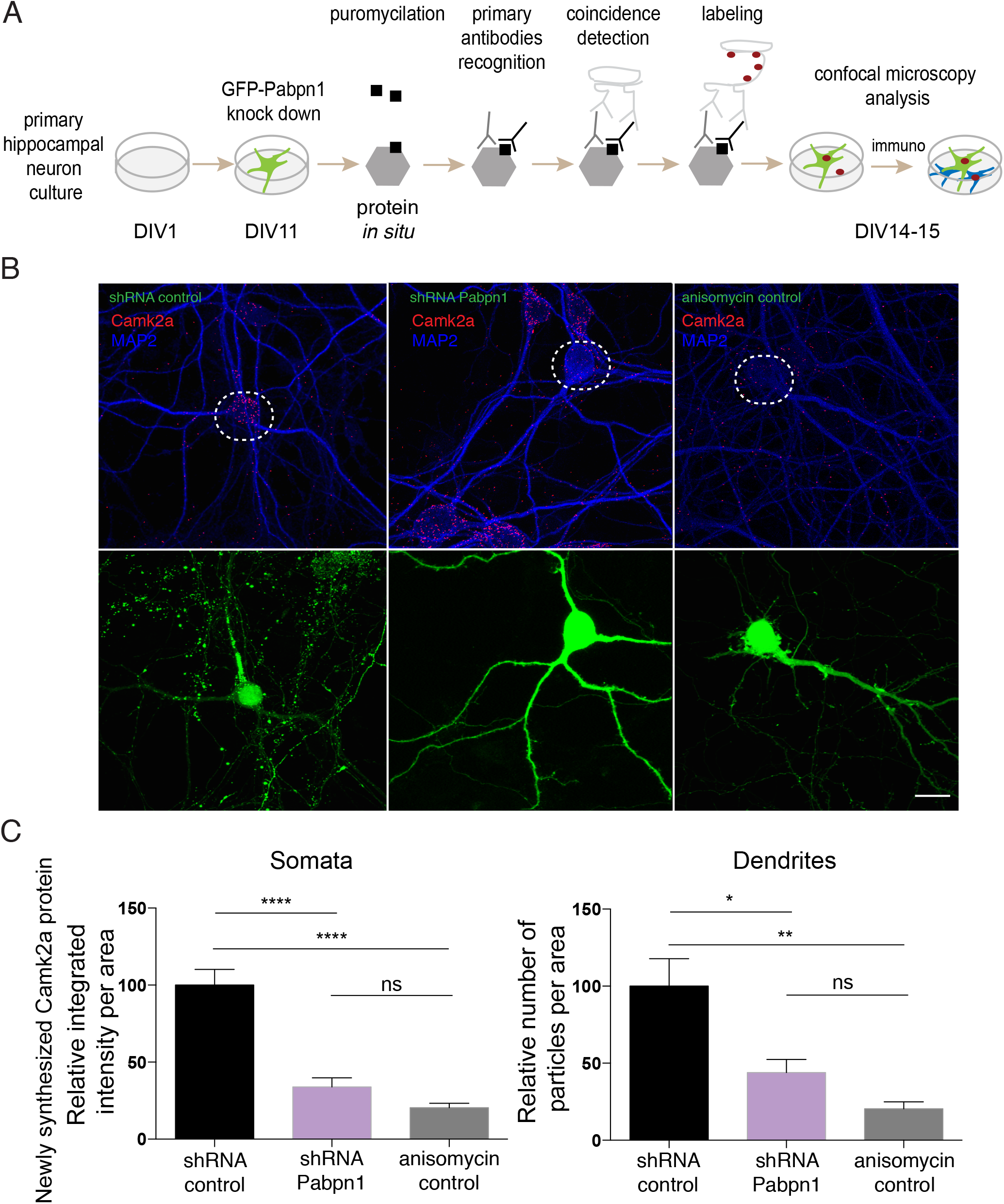
Newly synthesized Camk2a protein decreases upon Pabpn1 knock down. A. Schematic description of the puro-PLA method. DIV (days *in vitro*). B. Representative images of rat hippocampal culture upon shRNA control, shRNA against Pabpn1 and anisomycin control (GFP transfected neurons shown in bottom panel (green)). Newly synthesized Camk2a protein is highlighted in red and MAP2 stains neurons in blue. White dashed circle marks transfected somata. C. Quantification of newly synthesized Camk2a protein in somata and dendrites comparing the shRNA control, shRNA against Pabpn1 and anisomycin controls conditions. For somata n= 18, 13, 10 respectively in different groups, for dendrites n= 13, 9, 6. ****, p<0.0001; **, p=0.0059; *, p= 0.0289, non-significant (n.s.). Whiskers represent S.E.M.

### Pabpn1 knock down interferes with homeostatic plasticity

Given the above regulation of both Camk2a and Gria2, we next determined whether Pabpn1 and polyadenylation play a role in homeostatic plasticity. We elicited homeostatic plasticity using a 24 hours treatment of bicuculline a gamma-aminobutyric acid A (GABA_A_) receptor antagonist that elicits homeostatic down-scaling of miniature excitatory postsynaptic current (mEPSC) amplitudes (Turrigiano et al., 1998). In untreated or control shRNA neurons, bicuculline treatment resulted in a down-scaling of synaptic transmission (Figure 5A). In contrast, neurons that were exposed to shRNA against Pabpn1 failed to exhibit homeostatic scaling. To examine whether Pabpn1 itself may be regulated by the bicuculline treatment we measured Pabpn1 mRNA expression levels with *in situ* hybridization (using probes directed against the Pabpn1 coding sequence probe as well as qPCR (Figure 5B, C). In both cases we detected no change in Pabpn1 expression levels following BIC treatment, nor were there changes in Pabpn1 protein, as assessed by immunostaining (Figure 5D). Thus, the observed block of homeostatic plasticity following Pabpn1-KD is caused by effects on downstream plasticity genes that are targeted by Pabpn1 and potentially regulated by effect on polyA tail length and/or alternative polyadenylation.

**Figure 5.**
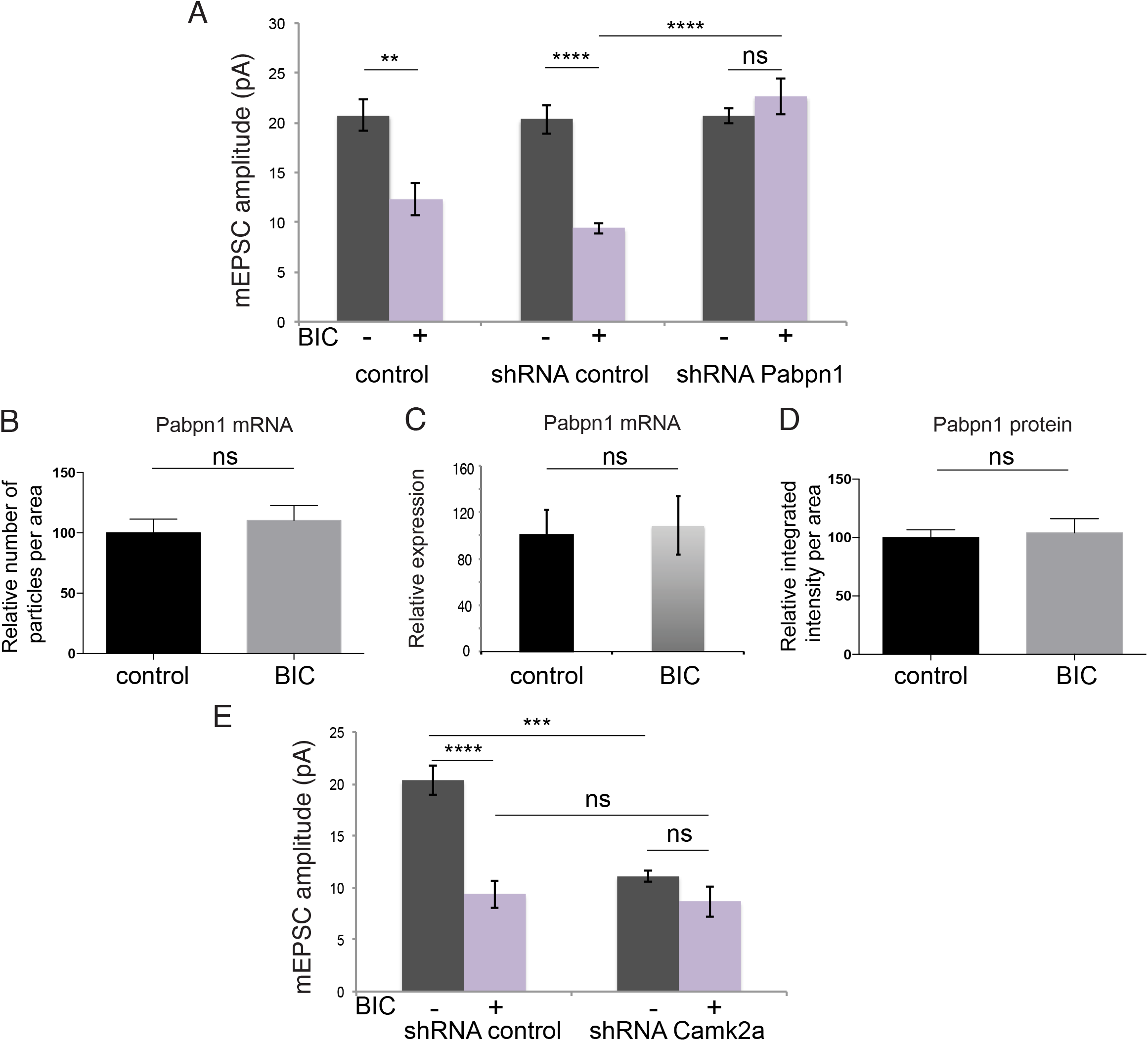
Pabpn1 knock-down blocks homeostatic plasticity. A. Mini excitatory postsynaptic current (mEPSC) amplitude measures are shown for the following conditions: control (untransfected cells: BIC (−), n=8; BIC (+), n=7), shRNA control (non-targeting control: BIC (−), n=8; BIC (+), n=7) and shRNA against Pabpn1 (BIC (−), n=8; BIC (+), n=8). BIC treatment led to decrease in amplitude in untransfected control and shRNA control conditions (**p=0.0019, ****p<0.0001 respectively). mEPSC amplitude did not change following BIC treatment of Pabpn1-KD cells (n.s.). A significant difference between the shRNA control BIC and shRNA against Pabpn1 BIC conditions was observed (****p<0.0001). B. Quantification of *in situ* hybridization against Pabpn1 mRNA w/o BIC treatment, show no difference between the control and BIC treated samples. C. Q-PCR data shows no difference in expression of Pabpn1 mRNA following BIC treatment. Three technical replicates are normalized to the housekeeping gene Rnr1. Whiskers represent standard deviation (S.D.). D. Quantification of the immunostaining of Pabpn1 in the control condition and following BIC treatment shows no significant effect (n.s.). Whiskers represent S.E.M. E. Mini excitatory postsynaptic current (mEPSC) amplitude measures are shown in shRNA control (BIC (−), n=8; BIC (+), n=7) and shRNA against Camk2a (BIC (−), n=8; BIC (+), n=8). BIC treated shRNA control decreased the mEPSCs amplitude significantly compared to untreated shRNA control neurons (****p<0.0001). shRNA against Camk2a showed a decrease in the mEPSCs amplitude (***p=0.0001) compared to shRNA control.

To directly test if a decrease in Camk2a protein elicited by Pabpn1-KD (Figure 2 and 3) causes the block of homeostatic scaling we knocked-down Camk2a using an shRNA approach. The treatment of neurons with an shRNA against Camk2a resulted in an 35% reduction in Camk2a protein, (data not shown). The KD of Camk2a protein resulted in a decrease in the mEPSC amplitude (Figure 5). The addition of bicuculline, however, did not elicit a further down-scaling of the mEPSC amplitude in the Camk2a-KD neurons (Figure 5E). There are two important aspects of this experiment to interpret. The first aspect concerns whether or not homeostatic scaling was affected by either KD, and in this case both KDs blocked scaling, consistent with the possibility that the effect of Pabpn1-KD on plasticity could be due to the reduction in Camk2a protein. The second consideration is the extent to which, under baseline conditions, the KD of Pabpn1 and Camk2a elicit similar phenotypes. The KD of Camk2a resulted in a reduction in the basal mEPSC amplitude which was not observed in the Pabpn1 KD. As the Pabpn1 KD resulted in a 56% reduction of Camk2a protein and the Camk2a KD resulted in a 35% percent reduction of Camk2a protein, this is unlikely to be due to differences in KD efficiency. Alternatively the difference could be due to the fact that Pabpn1 has multiple targets and a change in one or more of its other targets could compensate for the effect of Camk2a KD on the basal mEPSC amplitude.

## Discussion

Based on a model proposed by Jenal and colleagues (Jenal et al., 2012) we bioinformatically screened 3’end sequencing libraries of hippocampal neurons and predicted >1200 Pabpn1 targets. Using confocal microscopy we examined several key synaptic plasticity genes from the predicted targets dataset and found that Pabpn1 regulates polyadenylation of Camk2a and Gria2 but does not regulate Grin2a. Based on an observed decrease in both 3’UTR and coding sequence of Camk2a and Gria2, we suggest that the mechanism leading to the detected phenotype includes the regulation of polyA tail length and subsequent effects on the transcript stability and translation. The proposed mechanism is in line with our finding that Pabpn1-KD elicited a decrease in newly synthesized Camk2a protein.

In this study, we detected a significant downregulation of Camk2a in dendrites following Pabpn1-KD. Camk2a mRNA codes for the Calcium/Calmodulin Dependent Protein Kinase II Alpha (CamkIIα), a highly abundant protein in neurons and key mediator of synaptic plasticity (Miller et al., 2002; Wayman et al., 2008). Using mutant mice in which the Camk2a dendritic localization element was disrupted, Miller and colleagues demonstrated the importance of dendritically localized Camk2a for some forms of memory (Miller et al., 2002). Taking these findings into consideration, our data showing that Pabpn1-KD leads to a significant downregulation of Camk2a in dendrites, as well as a block of homeostatic plasticity suggests that the disruption of Camk2a function may be, at least in part, responsible for the block of homeostatic scaling following Pabpn1-KD. In contrast, the effect of Camk2a-KD on basal mEPSC amplitude was not phenocopied by Pabpn1-KD. This suggests that Pabpn1-KD has other effects that compensate for the reduction in mEPSC amplitude observed following Camk2a-KD. We suggest that changes in some of the predicted but not studied Pabpn1 targets may also regulate mEPSC amplitude.

The second examined key synaptic plasticity gene was Gria2 encoding the GluR2 subunit of the AMPAR. AMPARs mediate excitatory transmission in the brain. Changes in the number of AMPARs are crucial for homeostatic plasticity. Goold and Nicoll (2010) found that synaptic depression of AMPARs requires the GluR2 AMPAR subunit. The authors measured EPSCs following shRNA against GluR2 and detected a block of synaptic scaling (Goold and Nicoll, 2010). We would predict a decrease in the amplitude of basal mEPSCs upon GluR2-KD, similar to what we observed for Camk2a. Therefore, we suggest that change in GluR2 is also not the cause or the sole cause of the block of homeostatic scaling we observed following Pabpn1-KD.

Taken together, our study shows that Pabpn1 regulates polyadenylation of the key synaptic plasticity genes Camk2a and Gria2, while based on bioinformatic predictions it may also regulate multiple other targets in neurons. Further, Pabpn1 deficiency impairs homeostatic plasticity by a mechanism that we suggest consists of multiple molecular players including but not restricted to Camk2a and Gria2. In future studies, 3’end sequencing of Pabpn1-depleted cells in combination with RNA interference strategies against multiple targets may dissect this mechanism and further shed light on the roles of Pabpn1 and polyadenylation in homeostatic plasticity. Finally, our findings show a potential link between Pabpn1 and neurological disorders in which plasticity is impaired.

## Materials and methods

### Primary hippocampal neuron culture

Primary hippocampal neuronal cultures were prepared from hippocampi of day 1 Sprague Dawley male and female rat pups as described previously (Aakalu et al., 2001). Neurons were plated onto poly-D-lysine-coated glass-bottom MatTek dishes (density: 20 to 40 × 10^3^ cells/cm^2^) and maintained in Neurobasal A medium containing B-27 and glutamax supplements (Invitrogen) at 37°C until they were used (DIV 11-20). Experiments were performed according to the regulations of German animal welfare law.

### RNA interference experiments

Knock down of Pabpn1 and Camk2a were performed using SureSilencing shRNAs, Qiagen (GFP containing clones with sequences: Pabpn1 = ACAGCTCCAGATCTCGATTCT, Camk2a = GCCTGGACTTTCATCGATTCT). A non-targeting shRNA was used as a control (GFP containing clone with sequence: GGAATCTCATTCGATGCATAC). shRNA plasmids were amplified using One Shot^®^ TOP10 Competent Cells (Invitrogen) according to standard cloning protocols. Primary hippocampal neurons were transfected using a modified Magnetofectamin protocol, (Oz Biosciences), with 0.8 *µ*g of lipofectamine, 0.8 *µ*l of CombiMag and 0.8*µ*g of shRNA plasmid, incubated for 20-25 min and washed using NGM supplemented with glutamax, Invitrogen. The effectiveness of knock-down was screened after 3-4 days.

### Pharmacological treatments

Bicuculline treatments were performed by adding (-)-Bicuculline methochloride (Tocris, Cat. No. 0131, stock 100 mM in water, 40 *µ*M final) to the primary neuron culture medium for 12- 14 hours for imaging and RNA isolation experiments, and for 12- 20 hours for electrophysiology experiments. Anisomycin (Tocris, stock 100 mM in DMSO, 40 μM final) was preincubated with cells for 30 min on 37 °C in Puro-PLA control experiments. Puromycin (puromycin dihydrochloride, Sigma, stock 90 mM in water, 1μM final) was applied to all control and shRNA treated neurons at 37 °C for 4 min in Puro-PLA experiments.

### *In situ* hybridization and immunostaining in primary neurons

*In situ* hybridization was performed using the QuantiGene ViewRNA^™^ mRNA ISH Cell Assay with custom made probes targeting the 3’UTR long or coding sequence of the respective mRNAs. The probes were produced by Affymetrix recognizing Pabpn1 CDS (VC6-13823), Camk2a CDS (VC6-11639), Gria2 CDS (VC6-11359), Grin2a CDS (VC6-11368), Camk2a long UTR (VC1-14332), Gria2 long UTR (VC1-19395) and Grin2a long UTR (VC1-15554). In short, rat hippocampal neurons were fixed at DIV 14-21 for 30 min with a 4% paraformaldehyde solution (4% paraformaldehyde, 5.4% Glucose, 0.01M sodium metaperiodate in lysine-phosphate buffer). The *in situ* hybridization was performed following the manufacturer’s protocol omitting the protease treatment step. In hybridization step probes were used at a 1:50 dilution (for Pabpn1) or 1:75 dilution (all other probes). The amplification steps were performed using ViewRNA HC Screening Amplification Kits, Type 1 containing the label probe with an excitation at 547 nm or Type 6 containing label probe with an excitation at 647 nm at 1:100 dilutions of all components, as recommended. Following *in situ*, dendrites were immunostained using the following antibodies: ms-anti-Map2 (Sigma, M9942, 1:1000), rb-anti-Map2 (Millipore AB5622, 1:1000 dilution) or gp-anti-MAP2 (Synaptic Systems, 188004, 1:2,000) for 1h on room temperature. Secondary antibodies Alexa 405 anti corresponding species, all 1:1000, Life Technologies were applied for 30-45 minutes on room temperature. Cells were mounted or stored in 1x PBS and imaged. The experiments were performed in three biological replicates, each of them was prepared from a separate hippocampal neuron culture preparation.

For immunostainings of Pabpn1, neurons were fixed as described for *in situ hybridization*. After fixation, neurons were incubated for two times 10 min with 0.1 M glycin in blocking buffer (4% goat serum in PBS) and for 15 min with 0.5% triton in blocking buffer. Blocking was done for 1h, as the first antibody rb anti-Pabpn1, Abcam, ab75855, 1:250 was used for 1h and as the secondary antibody Alexa 546 (A11035) or Alexa 647 (A21245), Life Technologies, 1:1000 were used for 30-45 min. All steps were performed at room temperature. Neurons were washed and stored in 1x PBS prior to imaging.

### Immunostaining in tissues

Whole brain and hippocampi were isolated from adult wild-type rats and fixed in 4%PFA/PBS over night at 4 °C. Using a vibratome (Leica VT-1200), 50-100 μm thick slices were prepared. All subsequent steps for immunostaining were performed free-floating in 24 well-plates. Slices were permeabilized for 30 min at room temperature in 1%Triton/TBS and blocked for 1 hour in 1%BSA/1%Triton/TBS. Primary antibody staining was performed in blocking buffer overnight (rb anti-Pabpn1, Abcam, ab75855, 1:250 and ms-anti-Map2, Sigma, M9942, 1:1000). Slices were then washed three times in TBS, rocking for 10 min each time. Sections were incubated in secondary antibody/blocking buffer and 4,6-Diamidino-2-phenylindole, dihydrochloride (DAPI), 1:1000 for 2 hours at room temperature, washed three times, 10 min for each washing step. Slices were mounted and stored at 4 °C until imaging.

### Puro-PLA assay

Puro-PLA assay was performed as previously described (tom Dieck et al., 2015) with the following modifications. Briefly, cells were pretreated with anisomycin for 30 min and treated with puromycin for 4 min, or not treated as a control. Cells were fixed for 15 min, permeabilized for 10 min and blocked for 1h. Primary antibodies mouse anti-Puromycin, 1:3500, Kerafast, EQ0001 and rabbit anti-Camk2a, 1:1100, Millipore, 04-1079 were applied for 1.5 hour, washed and proximity ligation probes (PLA anti-rabbit PLUS, DUO90002 and PLA probe anti-mouse MINUS, DUO90004, both Duolink *in situ*, Sigma Aldrich) were applied for 1 hour at 37 °C. The ligation was performed for 30 min and the amplification for 100 min using the Duolink Detection Reagents Red DUO92008. Neurons were post-fixed for 10 min and immunostained following a standard protocol using the following antibodies: gp-anti-MAP2 (Synaptic Systems, 188004, 1:2,000) and donkey anti-gp Cy5, Dianova, 706-175-148, 1:500. Neurons were stored in 1x PBS until imaged.

### Image acquisition and processing

Images were acquired using a Zeiss LSM780 confocal laser fluorescence microscope system. Typically confocal images of 15-30 confocal planes were taken at optimal intervals using a 40x oil immersion objective. Somata of GFP positive neurons were circumscribed and dendrites were straightened using ImageJ software. Image analysis of *in situ* data from somata was performed using a custom MATLAB script where the average fluorescence intensity was measured per Z-stack and further normalized to the MAP2 area of the circumscribed cell body. In dendrites the particle number was determined using the same MATLAB script. The number of particles was normalized to the area of the straightened dendrite. Statistical analysis was performed in few steps. First, outliers were identified and removed by the ROUT method (Q=1%). Second, the data was tested for normality using D’Agostino-Pearson omnibus normality test and finally the data that exhibited a normal distribution was analyzed using unpaired *t*-test assuming both populations have the same standard deviation (S.D.). In cases where the data were not distributed normally, a Mann-Whitney-U nonparametric test was used.

To quantify Puro-PLA particles in somata, image analysis was performed using a custom script as previously described (tom Dieck et al., 2015). Particle analysis in dendrites, removal of outliers and normality tests were performed as already described for analysis of the *in situ* hybridization data. Statistical differences between groups were determined using Ordinary One-way ANOVA followed by Tukey’s multiple comparisons test.

Quantification of Pabpn1 immunostaining was performed using the Metamorph software. A low pass filter was applied to the images. Regions containing nuclei were drawn and the integrated intensity of each Z stack was calculated. Average integrated intensity by stack was calculated and divided by the area. Experiments are displayed as graphs after normalization of control to 100. Whiskers represent S.E.M.

### Quantitative PCR

Rat primary neurons grown at a density of 400 x10^3^ cells/cm^2^ were treated for 48 hours with 5 *µ*M cytosine β-D-arabinofuranoside (AraC), Sigma-Aldrich to deplete glia. Media was replaced and cells were further grown by adding 700 *µ*l of conditioned NGM media that contains glial products two times a week. On day 14, neurons were briefly washed with 1x PBS and scraped to TRIZOL (Thermo Fisher Scientific). RNA was extracted according to the standard TRIZOL protocol and treated with DNAseI for 30 min at 37 °C using the TURBO DNA-free kit (Thermo Fisher Scientific). Reverse transcription was performed using QuantiTect Reverse Transcription kit (Qiagen). Quantitative PCR was done using SYBR Green Master Mix, Roche and QuantiTect Assay Rn_Pabpn1_2_SG, QT02438744, Qiagen. Data was normalized to Rnr1 housekeeping gene (QuantiTect Assay Rn_Rnr1_1_SG, QT00199374, Qiagen).

### Electrophysiology

Rat hippocampal neurons transfected with shRNA against Pabpn1, shRNA against Camk2a or shRNA control were treated with 40 μM bicuculline in the culture media for 12- 20 hours while control cells were untreated or treated with bicuculline. Whole-cell patch-clamp recordings were performed on control and visually identified GFP-transfected neurons at room temperature. Signals were acquired at 50 KHz and Bessel filtered at 20 KHz using an Axopatch 200B amplifier. HEPES-buffered ACSF containing (in mM) 140 NaCl, 1.25 NaHPO_4_, 1 MgSO_4_, 3 KCl, 2 CaCl_2_, 1 MgCl_2_, 15 glucose, 10 HEPES [pH 7.4] was used as the bath solution. mEPSCs were recorded in the presence of 1 µM TTX and 20 µM bicuculline. The patch pipette internal solution contained (in mM) 120 potassium gluconate, 20 KCl, 0.1 EGTA, 2 MgCl_2_, 10 HEPES, 2 ATP, 0.4 GTP (pH 7.2, ∼300 Osm). Series resistance (R_s_, 6-20 MΩ) was not compensated but monitored in every sweep. R_s_ did not change more than ±3 MΩ in the cells included for analysis. Cells were voltage clamped at -70 mV. mEPSCs were analyzed offline using the Stimfit software. Minis were detected by template matching with 4 pA set as the amplitude threshold and the detected events were individually screened with a rise-time threshold set at <0.9 ms. Statistical differences between groups were determined by Ordinary one-way ANOVA with Tukey test corrected multiple comparisons.

### Bioinformatic analysis

3’end sequencing data were aligned using STAR software. mRNAs that possess the shorter isoform that ends with a less common poly(A) signal and extended 3’UTR isoform that ends with the canonical (AAUAAA) polyA site (PAS) were selected using a custom script.

## Author contributions

I. V. designed and performed the experiments and analysis, and wrote the manuscript. S. S. performed and analyzed the electrophysiology experiments. G. T. performed analysis. M.W. performed imunostainings in tissues. I. E., C. G., N. F helped with the experiments. I. C. performed the initial experiments. E.M.S. conceived and supervised the project, and wrote the manuscript.

## ACKNOWLEDGMENTS

We thank I. Bartnik, A. Staab, C. Thum and D. Vogel for the preparation of hippocampal neuron cultures. We thank M. Heumueller for using his custom made ImageJ based plugin for Puro-PLA particle analysis and L. Kochen for help with Puro-PLA. We also thank the students M. Azcorra, I. Uyan, G. Fachinger and C. Williams for their help in the optimization of the experiments and analysis. E.M.S. is funded by the Max Planck Society, an Advanced Investigator award from the European Research Council, DFG CRC 1080, DFG CRC 902, and the DFG Cluster of Excellence for Macromolecular Complexes.

## References

Aakalu, G., Smith, W.B., Nguyen, N., Jiang, C., and Schuman, E.M. (2001). Dynamic visualization of local protein synthesis in hippocampal neurons. Neuron 30, 489–502.

Beaulieu, Y.B., Kleinman, C.L., Landry-Voyer, A.M., Majewski, J., and Bachand, F. (2012). Polyadenylation-dependent control of long noncoding RNA expression by the poly(A)-binding protein nuclear 1. PLoS Genet 8, e1003078.

Brais, B., Bouchard, J.P., Xie, Y.G., Rochefort, D.L., Chretien, N., Tome, F.M., Lafreniere, R.G., Rommens, J.M., Uyama, E., Nohira, O., et al. (1998). Short GCG expansions in the PABP2 gene cause oculopharyngeal muscular dystrophy. Nat Genet 18, 164–167.

Bresson, S.M., and Conrad, N.K. (2013). The human nuclear poly(a)-binding protein promotes RNA hyperadenylation and decay. PLoS Genet 9, e1003893.

Chartier, A., Klein, P., Pierson, S., Barbezier, N., Gidaro, T., Casas, F., Carberry, S., Dowling, P., Maynadier, L., Bellec, M., et al. (2015). Mitochondrial dysfunction reveals the role of mRNA poly(A) tail regulation in oculopharyngeal muscular dystrophy pathogenesis. PLoS Genet 11, e1005092.

de Klerk, E., Venema, A., Anvar, S.Y., Goeman, J.J., Hu, O., Trollet, C., Dickson, G., den Dunnen, J.T., van der Maarel, S.M., Raz, V., et al. (2012). Poly(A) binding protein nuclear 1 levels affect alternative polyadenylation. Nucleic Acids Res 40, 9089–9101.

Derkach, V., Barria, A., and Soderling, T.R. (1999). Ca2+/calmodulin-kinase II enhances channel conductance of alpha-amino-3-hydroxy-5-methyl-4-isoxazolepropionate type glutamate receptors. Proceedings of the National Academy of Sciences of the United States of America 96, 3269–3274.

Elkon, R., Ugalde, A.P., and Agami, R. (2013). Alternative cleavage and polyadenylation: extent, regulation and function. Nat Rev Genet 14, 496–506.

Goold, C.P., and Nicoll, R.A. (2010). Single-cell optogenetic excitation drives homeostatic synaptic depression. Neuron 68, 512–528.

Hanson, P.I., Meyer, T., Stryer, L., and Schulman, H. (1994). Dual role of calmodulin in autophosphorylation of multifunctional CaM kinase may underlie decoding of calcium signals. Neuron 12, 943–956.

Holt, C.E., and Schuman, E.M. (2013). The central dogma decentralized: new perspectives on RNA function and local translation in neurons. Neuron 80, 648–657.

Jenal, M., Elkon, R., Loayza-Puch, F., van Haaften, G., Kuhn, U., Menzies, F.M., Oude Vrielink, J.A., Bos, A.J., Drost, J., Rooijers, K., et al. (2012). The poly(A)-binding protein nuclear 1 suppresses alternative cleavage and polyadenylation sites. Cell 149, 538–553.

Kerwitz, Y., Kuhn, U., Lilie, H., Knoth, A., Scheuermann, T., Friedrich, H., Schwarz, E., and Wahle, E. (2003). Stimulation of poly(A) polymerase through a direct interaction with the nuclear poly(A) binding protein allosterically regulated by RNA. EMBO J 22, 3705–3714.

Kuhn, U., Gundel, M., Knoth, A., Kerwitz, Y., Rudel, S., and Wahle, E. (2009). Poly(A) tail length is controlled by the nuclear poly(A)-binding protein regulating the interaction between poly(A) polymerase and the cleavage and polyadenylation specificity factor. J Biol Chem 284, 22803–22814.

Martin, K.C., and Ephrussi, A. (2009). mRNA localization: gene expression in the spatial dimension. Cell 136, 719–730.

Miller, S., Yasuda, M., Coats, J.K., Jones, Y., Martone, M.E., and Mayford, M. (2002). Disruption of dendritic translation of CaMKIIalpha impairs stabilization of synaptic plasticity and memory consolidation. Neuron 36, 507–519.

Miura, P., Shenker, S., Andreu-Agullo, C., Westholm, J.O., and Lai, E.C. (2013). Widespread and extensive lengthening of 3' UTRs in the mammalian brain. Genome research 23, 812–825.

Schanzenbacher, C.T., Sambandan, S., Langer, J.D., and Schuman, E.M. (2016). Nascent Proteome Remodeling following Homeostatic Scaling at Hippocampal Synapses. Neuron 92, 358–371.

Taliaferro, J.M., Vidaki, M., Oliveira, R., Olson, S., Zhan, L., Saxena, T., Wang, E.T., Graveley, B.R., Gertler, F.B., Swanson, M.S., et al. (2016). Distal Alternative Last Exons Localize mRNAs to Neural Projections. Mol Cell 61, 821–833.

Tavanez, J.P., Calado, P., Braga, J., Lafarga, M., and Carmo-Fonseca, M. (2005). In vivo aggregation properties of the nuclear poly(A)-binding protein PABPN1. RNA 11, 752–762.

tom Dieck, S., Kochen, L., Hanus, C., Heumuller, M., Bartnik, I., Nassim-Assir, B., Merk, K., Mosler, T., Garg, S., Bunse, S., et al. (2015). Direct visualization of newly synthesized target proteins in situ. Nature methods 12, 411–414.

Tomita, S., Adesnik, H., Sekiguchi, M., Zhang, W., Wada, K., Howe, J.R., Nicoll, R.A., and Bredt, D.S. (2005). Stargazin modulates AMPA receptor gating and trafficking by distinct domains. Nature 435, 1052–1058.

Turrigiano, G.G., Leslie, K.R., Desai, N.S., Rutherford, L.C., and Nelson, S.B. (1998). Activity-dependent scaling of quantal amplitude in neocortical neurons. Nature 391, 892–896.

Wayman, G.A., Lee, Y.S., Tokumitsu, H., Silva, A.J., and Soderling, T.R. (2008). Calmodulin-kinases: modulators of neuronal development and plasticity. Neuron 59, 914–931.

Weill, L., Belloc, E., Bava, F.A., and Mendez, R. (2012). Translational control by changes in poly(A) tail length: recycling mRNAs. Nat Struct Mol Biol 19, 577–585.

